# Sufficiency of ITD for (biased) human auditory azimuthal perception

**DOI:** 10.1101/2024.11.05.622135

**Authors:** Anna Carolina Pereira Fernandez Okamoto de Azevedo, Tomás Guimarães Pellegrino, José Luis Peña, Bóris Marin, Rodrigo Pavão

**Author notes:** Equal contribution (Anna Carolina Pereira Fernandez Okamoto de Azevedo, Tomás Guimarães Pellegrino).

## Abstract

Experiments on human auditory perception have shown that interaural time difference (ITD) is sufficient to generate spatial percepts, even though stimuli containing only the ITD cue are perceived as being emitted from inside the head instead of from external locations at specific azimuths. These experiments are thus interpreted as “lateralization” instead of “localization” tasks. In fact, lateralized spatial perception has been quantified using tasks in which participants have to report their estimates by selecting a putative location inside the head, or matching the perceived position to sounds with a given interaural level difference. Therefore, these estimates are made with respect to internal frames of reference, but it is unclear whether these percepts have any significance for the more ecological problem of locating an external sound source.

In order to investigate the link between internalized spatial percepts and sound localization, we designed a new task in which subjects are instructed to report externalized azimuthal location for sounds containing only ITD cues. Despite the mismatch between an internalized percept having to be reported as emanating from an external location, subjects were able to estimate azimuths consistently. Furthermore, normalized estimates were indistinguishable from those obtained using traditional lateralization tasks. Our results revealed a direct relationship between perceived azimuths and ITD, which deviates from that obtained from acoustical analysis of binaural recordings, revealing estimation biases. Intriguingly, these results indicate that externalized percepts are not required for the generation of azimuthal percepts.

## INTRODUCTION

Determining the position of sound sources in space is fundamental across species. Humans mainly employ Interaural Time Differences (ITDs) for estimating the azimuth of low frequency sounds (<1500 Hz), while Interaural Level Differences (ILDs) are used at higher frequencies, for which ITDs are undetectable and ambiguous (**Fig. 1**) (Blauert 1982; Middlebrooks & Green 1991; Jones et al 2015).

**FIG. 1.**
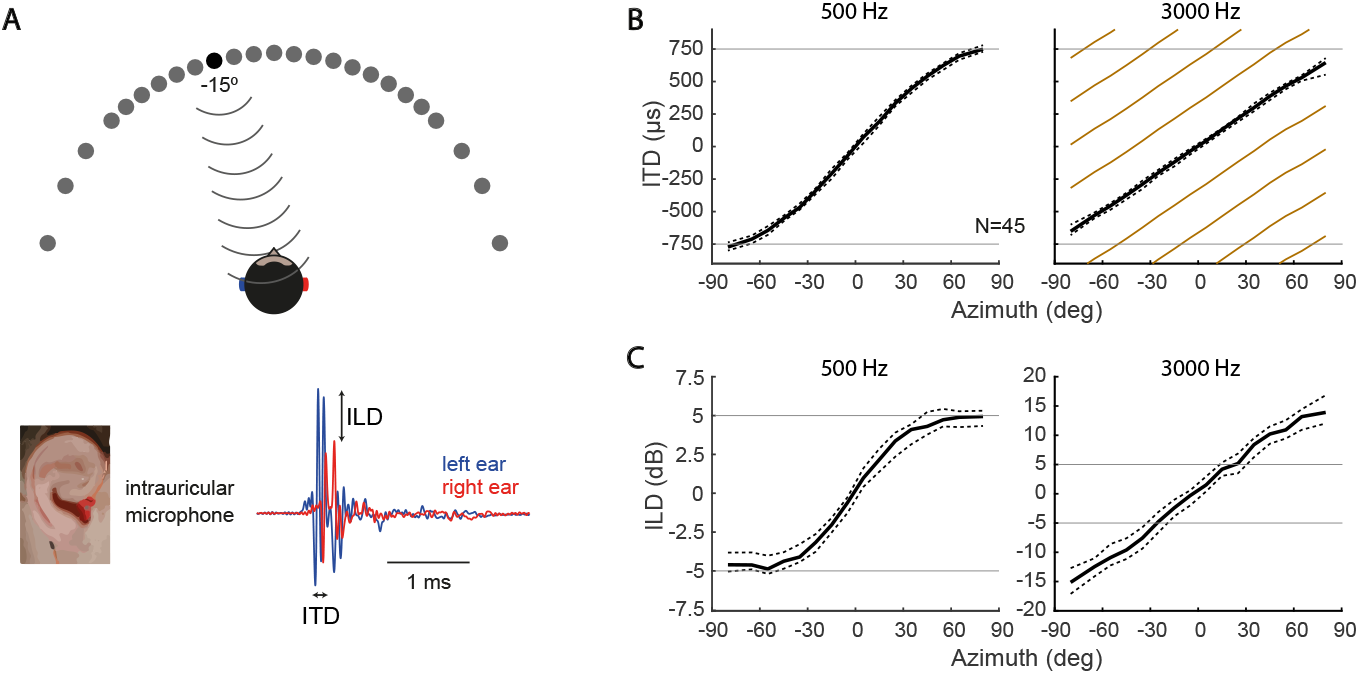
Acoustic cues used for sound localization in the horizontal plane. **(A)** Sound originates from point sources located at varying azimuths, reaching binaural microphones placed inside the subject’s ear canal. Below, recordings for a sound source at −15 degrees, and corresponding ITD and ILD cues computed from differences in timing and amplitude of the signal at each ear. **(B)** ITDs computed for sources at various azimuths for 500 Hz (left) and 3000 Hz (right) frequency components. Solid black lines indicate the median ITD across 45 subjects, while dashed lines indicate the interquartile range. This relationship is unique for 500 Hz, but ambiguous for 3000 Hz (beige lines). In fact, ITDs for components above 3000 Hz are not detected by humans, thus not used for sound localization. **(C)** ILD as a function of azimuth for the same frequency components as in B. Notice that the ILD range is narrower for 500 Hz (from -5 dB to 5 dB, saturated at the periphery) than for 3000 Hz (from -15 to 15 dB, without saturation). Analysis based on data extracted from the CIPIC database (Algazi et al. 2001).

Psychophysical studies of sound perception in the horizontal plane based on the ITD cue have traditionally employed *ILD-match* and *linear-bar* response methods (Yost 1981; Bernstein & Trahiotis 1985; Baumgartel & Dietz 2018). In both tasks, dichotic stimuli are presented through headphones. In the ILD-match method, the subject has to manipulate the ILD of a synthetic sound until it matches the perceived location of a sound stimulus lateralized by ITD (**Fig. 2A**). In the linear-bar method, the subject has to select a sound location by picking a point on a bar superimposed to a stylized human face (**Fig. 2B**). Notice that both tasks show saturated responses at extreme ITDs.

**FIG. 2.**
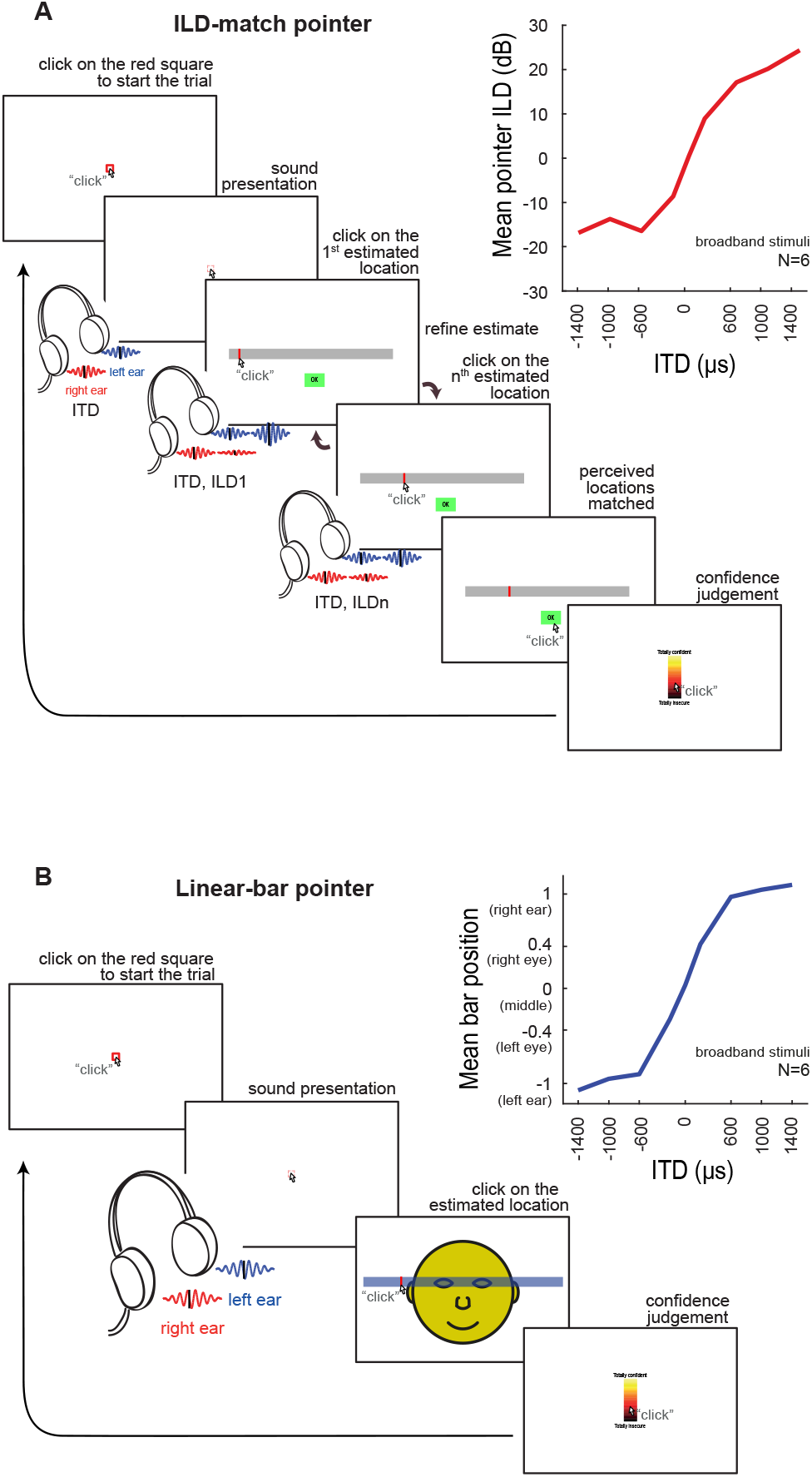
Traditional tasks for assessing lateralized perception of dichotic sounds varying on ITD. **(A)** Interface for the ILD-match task. The subject clicks inside the red square to start the trial (first screen), triggering the target ITD stimulus. The subject has then to pick a location in a horizontal bar, which will present a sound with an ILD corresponding to the selected position. This estimate can be further refined by selecting new points, until the subject feels that the position perceived from the original (ITD only) sound and the ILD based estimate match. Finally, the subject is asked to rate confidence in the estimation by selecting a point in a linear scale ranging from 0 to 100. Top right: ITD vs mean ILD responses of 6 subjects for broadband noise stimuli (adapted from Baumgartel, Dietz; 2018). **(B)** Interface for the linear-bar task. The procedure is similar to that described in A; there is, however, a single estimate, registered by clicking on a horizontal bar superimposed to a stylised human face. Top right: ITD vs mean position responses for broadband noise stimuli (also adapted from Baumgartel, Dietz; 2018).

Dichotic stimuli containing only ITD cues are perceived intracranially, i.e., as emanating from inside the head, and thus this spatial percept is referred to as sound “lateralization” instead of sound “localization” (Plenge, 1974), the latter applying to sounds originating in the environment. Both traditional tasks shown in **Fig. 2** are usually regarded as lateralization tasks and, as such, their response interfaces are not designed to collect azimuthal estimates. Nevertheless, slight adaptations of these interfaces should enable the extraction of azimuthal estimates, which could then be compared to those obtained from localization tasks, i.e. with sound sources placed in the environment. Such comparisons would be useful for investigating how ITD cues generate azimuthal percepts.

In the present study, we proposed the novel *azimuth pointer* task for testing whether azimuths can be estimated exclusively from ITD cues, and compared these estimates with those obtained through the traditional lateralization tasks. Results showed that the average and variability of azimuth estimates, as well as reaction times and confidence levels, are similar across tasks. This supports the hypothesis that ITD cues are sufficient for azimuth perception, even for internalized percepts. We also showed that the azimuth pointer task is as accurate as the traditional procedures for assessing spatial perception. Furthermore, our results indicate that azimuth estimation based solely on ITD cues deviate from the relationship between azimuth and ITD based solely on acoustic considerations: central ITD lead to estimates biased towards the periphery, while lateral ITD lead to estimates around a constant peripheral azimuth. Intriguingly, these results also suggest that externalization is not required for azimuth perception. Counterintuitively, these results also indicate that azimuth percepts can be generated even if the source is perceived to be inside the head.

## RESULTS

We collected lateralization estimates from 14 subjects using the *ILD-match* and *linear-bar* response methods (**Fig. 2**) (Baumgartel & Dietz 2018) as well as the novel *azimuth pointer* method (**Fig. 3A**), using the same dichotic stimuli: bursts of 500-Hz tones varying on ITD (**Fig. 3B**). For each of these tasks, we collected mean (**Fig. 3C** and **Fig. 4A**) and standard deviation (**Fig. 4B**) of the estimates, as well as reaction time and confidence level (**Fig. 4C**).

**FIG. 3.**
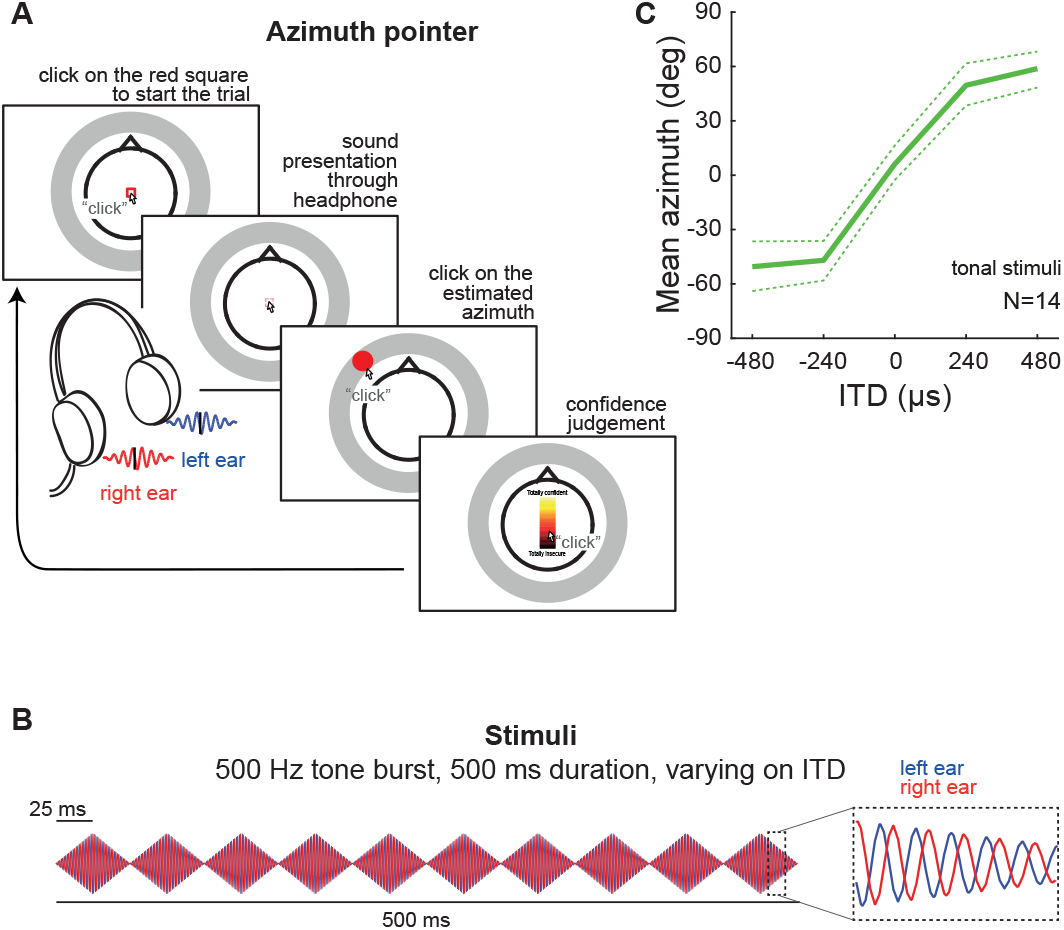
Azimuth pointer task for assessing azimuth perception of dichotic sounds varying on ITD. **(A)** Outline of each trial. The procedure is similar to the linear-bar task in Fig. 2B, but the subject is now instructed to estimate the azimuth of the sound source. Responses are registered by picking a point in a ring drawn around a stylized human head, viewed from the top. **(B)** Stimuli consisted of 500 Hz tone bursts with varying ITD. Notice that the traditional tasks reported on Baumgartel & Dietz (2018) employed broadband noise (Fig. 2). **(C)** Mean azimuth estimates. The solid line represents average across subjects; dashed lines represent its 95% confidence intervals.

**FIG. 4.**
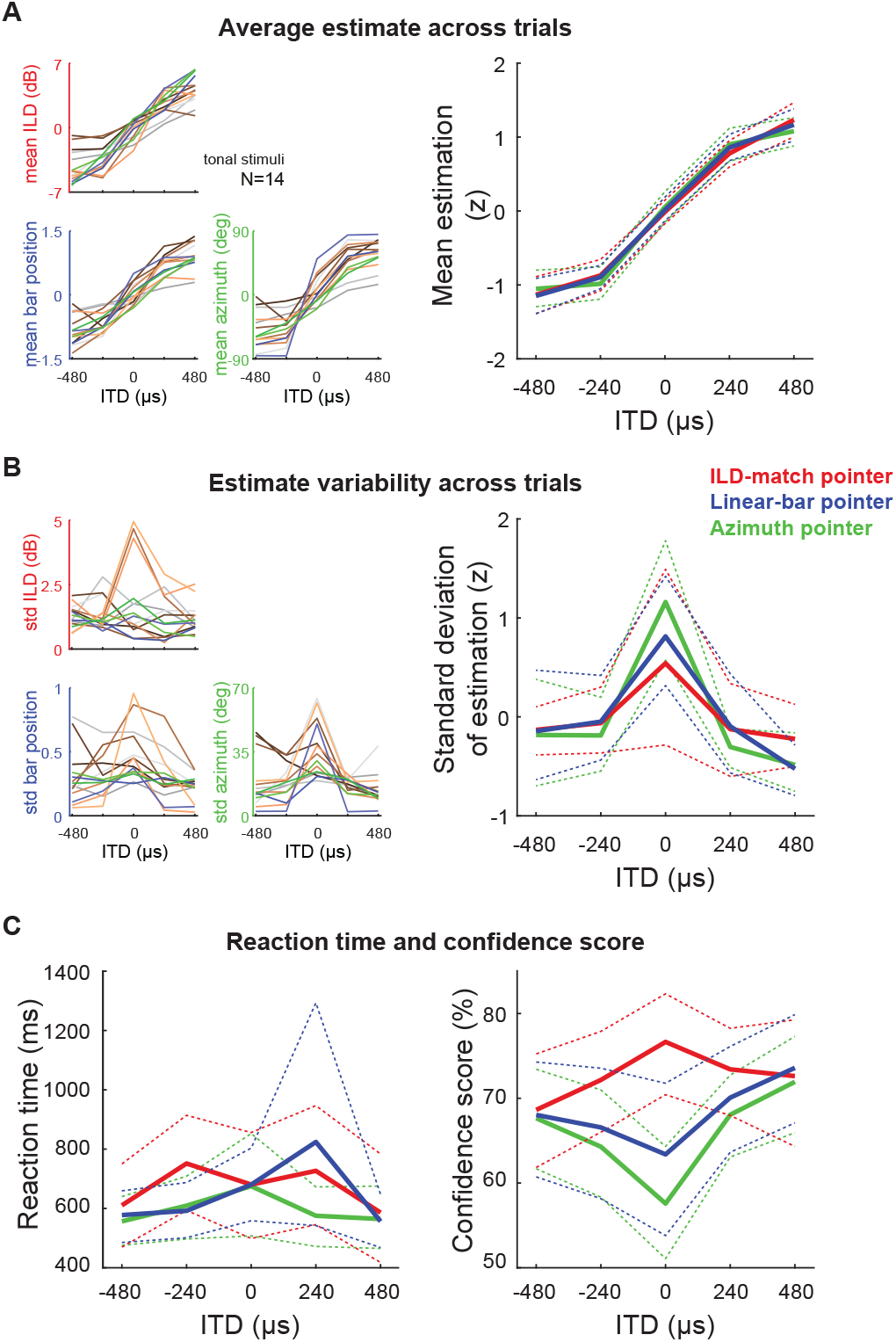
Comparing ITD-based spatial perception using the traditional and novel azimuth pointer tasks. **(A)** Mean estimation for the traditional (ILD-match and linear-bar) and new (azimuth pointer) tasks (left). Left: mean estimates for each of the three tasks across trials for each subject (color coded). Right: mean estimates averaged across subjects; the original data in distinct scales was normalized by computing z-scores (red, blue and green indicate ILD-match, linear-bar and azimuth pointer tasks, respectively). Note that the normalized mean estimates are almost indistinguishable across tasks. **(B)** Same as panel A, but for variability in the estimate across trials (right). Again, notice that the normalized standard deviations of estimates are similar across tasks. **(C)** Mean reaction time (left) and confidence in the accuracy of estimation for each task (right). Dashed lines represent the 95% confidence intervals around the means.

**Fig. 4A** shows mean and standard deviation of estimates for each of the three tasks. The mean estimates are strikingly similar across tasks, following the sigmoidal pattern seen in previous studies, saturating at peripheral ITDs (insets in **Fig. 2**; differences in dynamic range may be ascribed to the use of stimuli of distinct bandwidth). Mean azimuthal estimates averaged across subjects for the ILD-match, linear-bar and azimuth pointer tasks ranged from −4.2 to 4.2 dB, −0.8 to 0.9 (left and right ears correspond to −1 and 1), and −50.5 to 58.8 degrees, respectively. In order to compare estimates in different scales, we calculated z-scores for pooled subject responses.

Estimate variability across trials for each task exhibited the same pattern of higher variability at ITD 0 μs (front) in comparison to peripheral locations (**Fig. 4B**). Variability of estimates averaged across subjects for ILD-match, linear-bar and azimuth pointer tasks ranged 1.2 to 1.9 dB, 0.2 to 0.5 (considering that left to right ear indicates −1 to 1), and 14.0 to 36.3 degrees, respectively.

Fig. 4C shows reaction time and confidence scores collected using the same scale across response methods. Mean reaction times show large variability across subjects, but are in the same range across methods (**Fig. 4C**, left). Reaction time was defined as the interval between the onset of stimulus presentation and the start of the movement toward the response position. Given that there are multiple responses for the ILD-match task, we considered the interval between the first stimulus presentation and the start of the first response. Additionally, we compared the mean reaction times of the two traditional tasks to the azimuth pointer task (distance between green and red/blue curves in **Fig. 4C**). The ITD averaged reaction time for the linear-bar task was 50 ms longer than that of the azimuth pointer task; however, this difference did not reach statistical significance (p-value 0.26 in a bicaudal paired comparison). Consistently, the average first response time on the ILD-match was 75 ms longer than that for the azimuth pointer task, but once again the difference did not reach statistical significance (p-value 0.35).

The subjects’ confidence on the accuracy of their estimation was similar only between the linear-bar and azimuth pointer tasks: peripheral ITDs were estimated with higher confidence (**Fig. 4C**, right). The opposite pattern was found for the ILD-match task, in which frontal ITDs led to higher confidence scores. This is probably related to the fact that for each trial in this task, the subject is instructed to refine his estimate until the ILD based response matches the ITD stimulus, while the other tasks had only one presentation per trial. In the ILD-match task, subjects performed more iterations and reported lower confidence for peripheral (in comparison to frontal) stimuli.

**Fig. 5A** shows the acoustic mapping between azimuth and ITD (same as **Fig. 1B** but with inverted axes), calculated from the Head Related Impulse Responses (HRIR − as described in the Methods section), as well as the psychophysical mapping between stimulus ITD and estimated azimuth obtained from the azimuth pointer task. Notice that these curves have opposite concavities: the acoustic curve saturates at ±750 μs, while azimuth estimates saturate at ±50 degrees. These patterns indicate that azimuth perception from ITD cues is biased. We then obtained the azimuth perceptual bias across ITD by subtracting the acoustic curve from the psychophysics estimates (**Fig. 5B**). Estimates were maximally accurate for 0 μs ITD, maximally biased toward the periphery at ±230 μs (bias of around 30 degrees), and finally biased towards the front for more extreme ITDs.

**FIG. 5.**
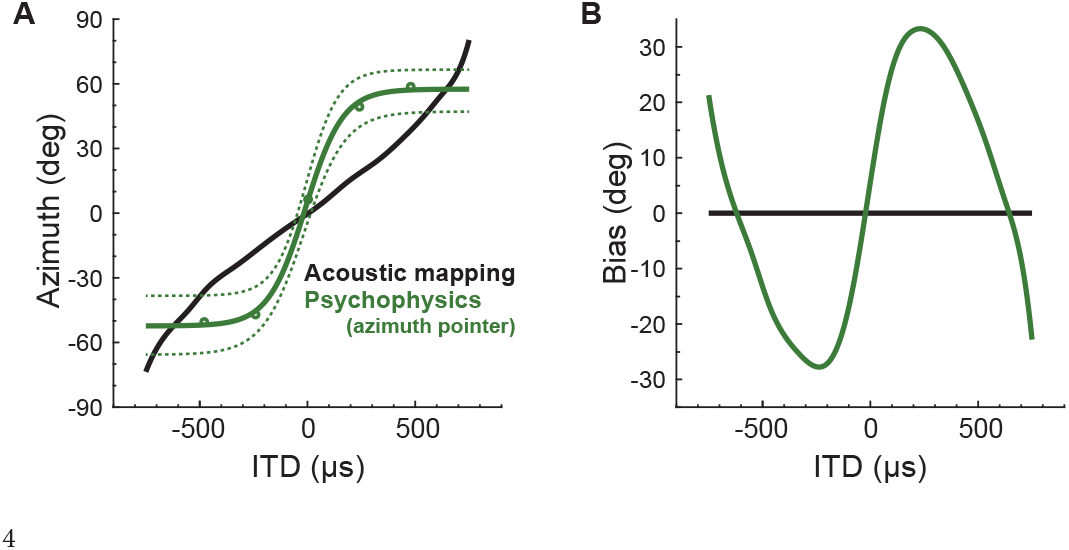
Azimuth estimation biases as a function of stimulus ITD. **(A)** Black curve: acoustic mapping between ITD and azimuth (same as Fig. 1B left) calculated from the HRIR. Green curve: sigmoid fit to the estimates obtained with the azimuth pointer task, for tone bursts with varying ITD. Dotted curves represent the 95% confidence interval of the mean. **(B)** Biases in azimuth estimation (green curve) highlighted by subtracting the acoustic relationship from the estimates.

## DISCUSSION

### Localization vs lateralization: is externalization necessary for azimuth perception?

Sounds presented through binaural headphones and containing only ITD cues are perceived as being lateralized, as opposed to emanating from an external location in the environment (Plenge, 1974). Traditional tasks for evaluating this percept (ILD-matching and linear-bar tasks) enforce lateralized responses (Yost 1981; Bernstein & Trahiotis 1991; Baumgartel & Dietz 2018). Nevertheless, we have shown that it is possible to consistently estimate azimuth from stimuli containing only ITD cues, using our novel azimuth pointer task (**Fig. 3**).

If azimuth estimates were constructed from the internal (lateralized) percepts as an additional computational step along the auditory pathway, that would be reflected in increased reaction times for the azimuth pointer task in comparison to the traditional tasks. But that is not the case: reaction times were similar across all response methods (**Fig. 4C**, left). Additionally, z-score normalized responses for each of the three methods were very similar, both on average and on variability across trials (**Fig. 4AB**, right), indicating that there is no tradeoff between reaction time and estimation accuracy/precision.

These results support the idea that externalization is not actually required for azimuthal perception. Even though it is possible to construct externalized dichotic stimuli by taking into account the acoustic transformations produced by head and room (Leclève et al 2019), cues that are critical for externalization (mainly echoes) carry little directional information (Wallach et al. 1949; Brown et al. 2015). These results motivate further investigations, in order to dissect the role of each cue (ITD, ILD, reverberation, spectra) and their interactions on externalization and the generation of azimuth percepts.

### Estimate saturation for peripheral ITDs and azimuthal perception biases

Baumgartel (2018) identified two issues with the ILD-match task: large inter-subject variability and influence of stimulus bandwidth. We argue that these problems stem from the ILD cue, which has larger variability and higher frequency dependence in comparison to the ITD cue (**Fig. 1BC**). We avoided these problems by normalizing the scale of responses across subjects and fixing all stimuli to 500 Hz, obtaining similar results for all three methods (**Fig. 4A**). Baumgartel (2018) also identified that the linear-bar task led to responses saturating at the periphery − which we also observed for the three tasks. Therefore, such perceptual biases seem to be a feature of spatial auditory perception from ITD cues (**Fig. 5**), rather than being a methodological artifact.

Our novel task not only allows extracting additional information from widely used psychophysical tasks with minor changes to the protocols, but also provides a systematic way to investigate the biases on ITD-based azimuth perception. We speculate that such biases may reflect the perceptual thresholds for detecting ITD changes, which vary across azimuth (Mills 1958, Brugguera 2013, Pavao et al 2020, Camperos et al 2022): for ITDs corresponding to peripheral stimuli, low ITD discriminability will lead to responses that saturate around a given azimuth (around ±50 deg in **Fig. 5A**). Consider, for example, ITDs around 650 μs (where the green and black curves intercept on **Fig. 5**). All estimates will cluster around 50 deg due to low discriminability, but for ITD lower than 650 μs there will be a bias towards the periphery, while ITDs higher than 650 μs will lead to a bias towards the center (**Fig. 5B**). On the other hand, high discriminability for frontal stimuli will lead to azimuth overestimation for ITD around 0 μs: small positive (negative) ITD will lead to estimates further to the right (left), leading to a bias towards the periphery for central ITD (**Fig. 5B**).

Thus, the bell-shaped zero-centered discriminability profile (reported in Pavao et al 2020 and Camperos et al 2022) is closely related to the slope of the response curve: high discriminability for low ITD values corresponds to positive slope, leading to estimates biased towards the periphery around zero; whereas low discriminability will lead to slopes close to zero, reflected as alternating center / periphery biases for extreme ITD values.

Interestingly, azimuthal estimation for free-field noise bursts (containing all spatial cues) lead to minimal saturation at the periphery (Yost et al 2013). Moreover, saturation ranges also vary with stimulus bandwidth: broadband noise lead to saturation of estimates around ±600 μs (**Fig. 2**, with data from Baumgartel, Dietz; 2018) while pure tones lead to saturation around ±300 μs for both traditional tasks (**Fig. 4a**). These results indicate that perceptual biases in estimation may reflect the amount of azimuthal information carried by each auditory spatial cue.

Taken together, these results indicate that the reductionist/unrealistic stimuli usually employed in psychophysical experiments lead to estimation biases (and internalized percepts) that can differ significantly from those for estimates based on naturalistic stimuli, which contain multiple interacting cues and exhibit statistical properties similar to the sounds animals encounter in their natural environments, under which their auditory systems have evolved.

## MATERIALS AND METHODS

### HRTF MEASUREMENTS

We estimated the acoustic relationship between azimuth and interaural cues (ITD and ILD) from head-related impulse responses (HRIR) of 45 subjects (Fig. 1) publicly available in the CIPIC HRTF Database (CIPIC Interface Laboratory at U.C. Davis, Algazi et al. 2001). Measurements were performed using loudspeakers mounted at various positions along a hoop, maintaining 1 m radius centered at the subject’s interaural axis. The subjects wore binaural microphones, blocking their ear canals.

### ITD and ILD analysis

In order to determine the acoustic mapping between ITD/ILD and azimuth, we started by convolving white noise signals with HRIRs from the CIPIC database, which were then filtered on frequency bands using a cochlear model (gamma-tone filter bank from Malcolm Slaney’s Auditory Toolbox, available at https://engineering.purdue.edu/~malcolm/interval/1998-010). Next, instantaneous phase and amplitude were extracted using the Hilbert transform (Matlab, Signal Processing Toolbox; Mathworks).

For each azimuth and cochlear band frequency *f*, we then computed the instantaneous interaural phase difference (IPD) from the instantaneous phase at each ear. Finally, we computed the circular mean IPD (radians) and transformed it to ITD (μs) according to 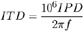, leading to the ITD-azimuth mapping for a single subject. Median and interquartile range were computed across subjects (Fig. 1B). In humans, the ITD-azimuth mapping is ambiguous for frequencies above 600 Hz, as reflected in the beige lines in Fig. 1B. These were obtained by adding and subtracting multiples of 2π to the mean IPD curve.

Following a similar procedure, we computed the instantaneous ILD from the signal amplitude *A* at each ear, using 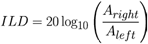. This was repeated for each cochlear filter frequency, azimuth and subject. Finally, we computed median ILD and its interquartile range across subjects (Fig. 1C).

### PSYCHOACOUSTIC TESTING

#### Subjects

We recruited 14 healthy participants (6 females and 8 males, age 27 ± 10, all right-handed). All participants were previously informed about the procedure before consenting to perform it. The protocol was approved by the Ethics committee of Universidade Federal do ABC, where the study took place.

#### ITD stimuli

For each trial, we presented bursts of 500-Hz tones through headphones (Pulse Max Motorola SH004), calibrated to 65 dBA on both sides (Instrutherm DEC-460 sound level meter). Based on a pilot study sampling multiple ITDs across the physiological range, we defined the critical stimulus range: five ITDs varying from -480 to 480 μs, in 240 μs steps (which corresponds roughly to azimuths -38, -17, 0, 17 and 38 degrees). Sound bursts consisted of ten consecutive 50-ms elements modulated with 25 ms rise and 25 ms fall time, and a total duration of 500 ms (**Fig. 3B**). Each participant performed the three estimation tasks distributed along 6 experimental blocks, randomly ordered for each subject. In each of the experimental blocks, every ITD was presented 9 times, totaling 45 trials per block; the first-order transitions between ITDs were controlled for avoiding sequence effects (Remillard 2008).

#### ILD-match task

We adapted the task of Bernstein & Trahiotis (1985) for using tonal burst stimuli and a slightly different trial structure (**Fig. 2A**). Subjects initiated trials by clicking on a red square on the computer screen, triggering a stimulus containing the target ITD cue. Next, the subject had to pick a point inside a horizontal bar drawn at the screen, triggering a stimulus containing the target ITD cue followed by a 500-Hz tone burst with ITD 0 μs and varying ILD, controlled by the point picked by the user (within the -7 to +7 dB range). Subjects could keep selecting points on the bar as many times as they wanted, until the perceived location of the user controlled ILD stimulus matched that of the target ITD stimulus.

#### Linear-bar task

We adapted the task of Yost (1981), including the same horizontal bar overlaid to an stylized head (frontal view, as in a mirror seen by the subject) as applied in Baumgartel & Dietz (2018) (**Fig. 2B**). The trials initiated when the subject clicked on a red square, triggering a stimulus containing the ITD cue. The subject then made his estimate by picking a point on the horizontal bar corresponding to the perceived location of the stimulus.

#### Azimuth pointer task

To test the hypothesis that azimuth perception can be induced by ITD stimuli, we adapted the linear-bar task by using a front instead of a top view stylised head, and replacing the horizontal bar by a ring around the head (**Fig. 3A**). Trials initiated when subjects clicked on a red square, triggering a stimulus containing the ITD cue. Next, the subject entered his estimate by picking a point on the ring, corresponding to the perceived azimuth.

#### Reaction time and confidence score

We registered reaction times for each of the three tasks by counting how long it took for the mouse pointer to leave the red square after the initial mouse click. Additionally, we collected a confidence score at the end of each trial, asking the subject to rate their estimation on a scale from 0 to 100%. The complete experiment lasted about 30 minutes, including breaks between blocks.

#### Data analysis

For every subject, we computed four summary statistics across the five ITDs: (1) mean of estimates, (2) standard deviation of estimates, (3) mean reaction time, and (4) mean confidence score. For each task, the mean and standard deviation of estimates were pooled across subjects then normalized: the distribution of 5 ITDs of each of the 14 subjects in the original scale of each task was transformed to the z scale. Next, we computed the mean across subjects and its 95% confidence interval (using bootstrap), as shown in **Fig. 4AB**. Mean reaction time and mean confidence were processed without normalization: we directly computed the mean across subjects as well as its 95% confidence interval (**Fig. 4C**). For comparing mean reaction times (averaged across ITDs) between pairs of tasks, we applied bootstrap procedures analogous to the paired t-test. Finally, for comparing the acoustic and psychophysical relationships between ITD and azimuth, we inverted the curve shown in **Fig. 1B**-left and computed the mean azimuth estimate across subjects and its 95% confidence interval (bootstrap). In order to map ITD to azimuth, we (1) interpolated the measurements using cubic splines for the acoustic relationship and (2) adjusted a sigmoid function to the azimuth estimates. All analyses were performed using Matlab, with built-in or custom-made routines.

## ACKNOWLEDGEMENTS

This work was supported by Coordenação de Aperfeiçoamento de Pessoal de Nível Superior (CAPES), Conselho Nacional de Desenvolvimento Científico e Tecnológico (CNPq) and National Institutes of Health (NIH grant DC007690).

## Notes

### Competing Interest Statement

The authors have declared no competing interest.

